# Fluorescence-lifetime optical electrophysiology in contracting cardiomyocytes

**DOI:** 10.64898/2026.01.12.699013

**Authors:** Euan Millar, Vytautas Zickus, Eline Huethorst, Giedrė Astrauskaitė, Jiuxuan Zhao, Gregor G. Taylor, Claudio Bruschini, Edoardo Charbon, Godfrey L. Smith, Caroline Müllenbroich, Daniele Faccio

## Abstract

Precise monitoring of cardiac electrophysiology in vitro is crucial to understanding heart function and cardiac disease. However, high-throughput, contact-free methods for directly measuring excitation-contraction coupling remain limited. Here, we introduce a new paradigm for quantitative electrophysiological imaging that combines fluorescence lifetime and intensity information to capture dynamic cardiac signals with high fidelity. We show that lifetime measurements are intrinsically decoupled from motion artifacts and provide calibrated calcium concentration and membrane potential readouts across wide fields of view. Using a gated single-photon avalanche diode camera, we acquire fluorescence lifetime images at up to 200 frames per second with sufficient signal-to-noise ratio such that each frame contains meaningful lifetime information without temporal averaging. This approach yields spatially resolved maps of absolute voltage and calcium values across contracting cardiomyocyte monolayers, revealing heterogeneous cell behaviors within individual assays and uncovering previously unreported dynamics during late-phase repolarization for real-time analysis of excitation-contraction coupling.

## INTRODUCTION

Cardiovascular diseases (CVDs) remain the leading cause of mortality, accounting for 13% of deaths world-wide [1]. Understanding cardiac electrophysiology at the cellular level under both healthy and pathological conditions is essential to uncovering the underlying mechanisms of CVDs and developing targeted treatments. In particular, high-resolution imaging of excitation-contraction coupling (voltage, calcium, and contractile dynamics, Fig. 1a) over a suitably large field of view (FOV) is necessary to resolve both cell-to-cell and regional differences in action potential (AP) and calcium transients within a syncytium of cardiomyocytes (CMs).

**FIG. 1.**
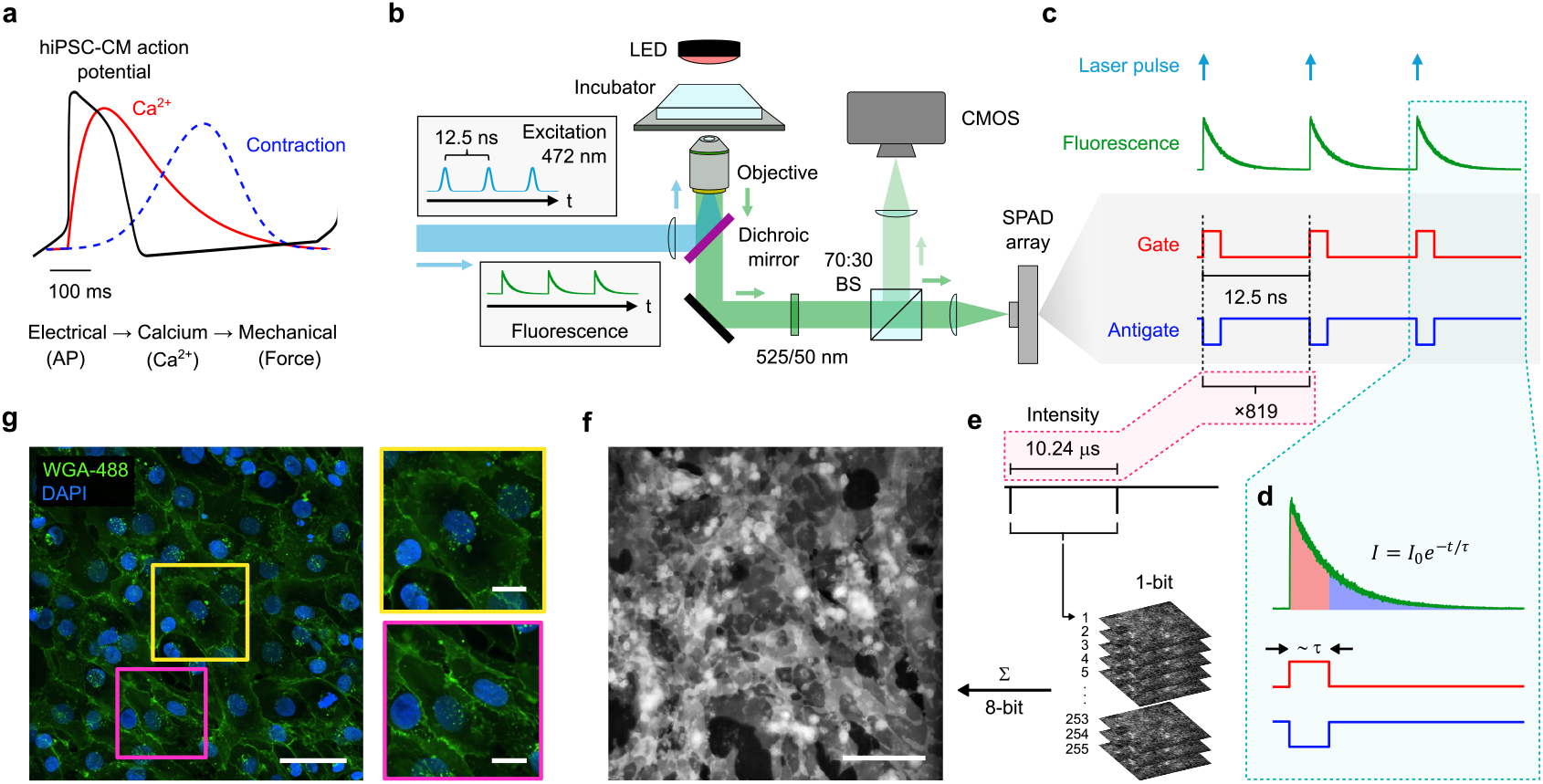
Overview of cardiac electrophysiology and fluorescence lifetime imaging system. (**a**) Diagram showing the excitation-contraction coupling mechanism in cardiac cells. Adapted from [18]. (**b**) Schematic of fluorescence lifetime imaging microscopy experimental setup. (**c**) Timing diagram showing the gated operation of the SPAD camera. The antigate profile is obtained by subtracting the gated images from the intensity frames. (**d**) Magnified view showing the gated capture of a single fluorescence emission. Shaded regions under the green curve highlight the regions integrated by each SPAD channel. (**e**) Multiple binary (1-bit) SPAD frames are accumulated and summed to form an 8-bit intensity image. Each binary exposure of 10.24 µs represents the total fluorescence signal integrated across 819 laser excitations. (**f** ) Example 8-bit fluorescence intensity image of a hiPSC-CM monolayer captured by the SPAD. White scale bar, 100 µm. (**g**) Immunofluorescence image of hiPSC-CM monolayer stained for Wheat-Germ Agglutinin-Alexa Fluor 488 (membranes) and DAPI (nuclei). White scale bar, 50 µm. Insets show magnified views of two regions indicated in the main panel. White scale bar, 15 µm.

Human-induced pluripotent stem cell-derived CMs (hiPSC-CMs) provide an important platform for such studies, with the ability to model human heart function [2, 3], provide insight into patient- and disease-specific drug responses [4], and enable the advancement of regenerative and personalized medicine efforts [5– 7]. However, current electrophysiology techniques are limited in their ability to capture these dynamic processes in live, contracting cells. For example, patch clamp recordings provide precise high-resolution voltage and ion channel data at the single-cell level but are invasive and low-throughput [8, 9]. Physical disruption of the cell membrane, the potential for leak currents, and the inability to record multicellular networks limit their use [10–12]. Multi-electrode arrays (MEAs) in contrast, allow non-invasive long-term recordings (up to weeks) from cell populations, but cannot resolve activity at the single-cell level or within subregions of a monolayer [8, 13]. MEAs also typically require contraction uncouplers, such as blebbistatin or 2,3-butanedione monoxime (BDM) [14– to reduce motion artifacts, which can alter electrophysiological parameters and confound results [17].

Optical methods that use voltage- and calcium-sensitive fluorescent dyes have enabled less invasive recordings with higher throughput [19– This approach infers changes in membrane potential (V_m_) and intracellular calcium ion (Ca^2+^) levels by measuring changes in fluorescence intensity at high imaging rates of ≥ 200 frames per second (fps) to ensure capture of the AP upstroke (5-10 ms). However, quantitative analysis is challenging due to confounding factors such as photo-bleaching, nonuniform dye distribution, and signal-to-noise limitations [19]. As a result, it is technically challenging and often impractical to extract values of V_m_ or Ca^2+^ concentrations from recorded traces of contracting cells. Like MEAs, fluorescence intensity measurements are also sensitive to motion [24]. Beyond motion uncouplers, ratiometric dyes can be used to overcome the challenges caused by motion artifacts. Others have used computational techniques to remove motion artifacts from untreated samples, but their efficacy is limited when processing images with high noise or reduced spatial resolution [25].

Fluorescence lifetime imaging microscopy (FLIM) addresses many of these limitations by offering a quantitative readout of absolute membrane potential and intracellular calcium levels through measurement of the excited-state lifetimes of fluorophores [26– This fluorescence lifetime measurement depends only on the temporal decay of the fluorescence signal rather than its absolute intensity, and is sensitive to the local environment of the fluorophore. However, current FLIM systems lack the high-speed sampling necessary to resolve AP spiking and other rapid cellular dynamics that occur within CM syncytiums [30]. Commercially available FLIM systems currently peak at 30 fps with 24-kilopixel resolution [31], dropping to ∼ 1 fps at 0.25 megapixels [32]. Although high-speed FLIM methods using streak cameras [33] or complex specialized setups [34] have captured electrical activity, they rely on expensive hardware, sacrifice resolution, and require averaging across multiple firing events, preventing accurate imaging of evolving intercellular phenomena. Gated cameras have also been applied to single molecule FLIM [35] and imaging at several frames/second at macro-to-microscopic scales of dynamical systems [36]. Here, we introduce fluorescence-lifetime optical electrophysiology (FLOE), a FLIM-enabled technique optimized for high-speed, high-resolution, quantitative measurements of voltage and calcium in hiPSC-CMs and other cellular preparations. Using a large-format, 500 × 500-pixel single-photon avalanche diode (SPAD) array with dual-gate acquisition [37], FLOE allows rapid life-time determination (RLD) [36] at rates of up to 200 fps – sufficient to resolve fast electrophysiological events such as the AP upstroke. Unlike conventional FLIM systems, FLOE provides complete lifetime information in a single frame, allowing continuous quantitative measurements of dynamic cellular processes. Critically, FLOE is insensitive to motion artifacts, removing the need for pharmacological uncoupling, and gives access to spatially-resolved, quantitative measurements of V_m_ and intracellular Ca^2+^ concentrations in beating cells. These capabilities open new avenues for studying cardiac disease mechanisms, drug responses, and intercellular dynamics in physiologically relevant models.

## RESULTS

### High-frame-rate FLIM

Our goal here is to quantitatively measure membrane potential and intracellular calcium dynamics while remaining insensitive to motion artifacts caused by contraction. To achieve this, we used a large-format SPAD array to perform high-frame-rate FLIM on cardiac cells during natural contractile behavior. A schematic of the microscope setup is shown in Fig. 1b and the gated operation of the SPAD is depicted in Fig. 1c. Synchronizing the camera with the excitation source, as seen in Fig. 1d, allows one to ‘gate’ the detection to correspond to distinct phases of fluorescence decay. This temporal gating allows for the discrimination of fluorescence lifetimes in a manner that is both rapid and sensitive (see Methods for details). Briefly, the system comprises an inverted fluorescence microscope with excitation provided by a 472 nm, 80 MHz pulsed laser diode. A red LED that is arranged above the objective provides bright-field imaging capability which, in combination with a CMOS camera, facilitates contractility measurements. Fluorescence intensity and lifetime measurements are performed using SwissSPAD3 [37], which operates at an 8-bit acquisition rate of 192 fps. This sensor performs synchronous dual-modality acquisition (Fig. 1c, e), capturing a time-integrated and time-gated image per exposure, with a minimum gate width of 1 ns. By default, the sensor also produces an anti-gate image as the difference between these two channels. Time-integrated intensity images (Fig. 1e) are formed by capturing 255 binary frames and summing them on the FPGA to form a single 8-bit image (Fig. 1f). This on-chip processing accounts for the discrepancy between the 10.24 µs binary exposure time and the 192 fps frame rate.

We apply this system to the measurement of excitation-contraction coupling in hiPSC-CM monolayer preparations similar to those shown in Fig. 1g. The dual-modality design of FLOE, capturing both time-integrated intensity and time-resolved lifetime information, enables a comprehensive characterization of dynamic fluorescence signals. By combining these complementary modalities, FLOE maximizes the strengths of each measurement, providing richer more reliable insights than either approach alone.

### Robust action potential imaging in contracting hiPSC-CM monolayers via FLIM

In intensity measurements, electrical activity is often reported as the instantaneous change in fluorescence intensity, Δ*F*, relative to a resting baseline fluorescence, *F*_0_ – denoted as Δ*F/F*_0_. However, any movement (such as CM contraction) directly alters the pixel-wise brightness of the measured fluorescence, causing fluctuations in Δ*F/F*_0_, i.e., motion artifacts. The solution we rely on is offered by certain dyes (such as FluoVolt) that also exhibit a voltage-dependent change in their excited-state lifetime that can track fluctuations in V_m_ [27]. Changes in fluorescence lifetime, *τ*, can be similarly expressed as Δ*τ/τ*_0_. Importantly, fluorescence lifetimes within specific cellular compartments tend to show strong spatial uniformity. Since the lifetime is an intrinsic property of a fluorophore and its environment and is thus independent of excitation intensity and local concentration, even if motion causes lateral shifts between spatial regions, the lifetime measurements are minimally perturbed. To assess this statement and our system’s ability to resolve individual APs under natural contractile conditions, we compared fluorescence intensity and fluorescence lifetime recordings in hiPSC-CM monolayers stained with FluoVolt before and after treatment with BDM.

Quantitative analysis of contraction dynamics before and after BDM treatment were performed using an open-source toolbox [38] and confirmed that BDM treatment reduced the mechanical contraction amplitude by 76%, see Fig. 2a and f. A comprehensive analysis of the change in contraction parameters is included in the Supplementary Material (Supplementary Fig. 1).

**FIG. 2.**
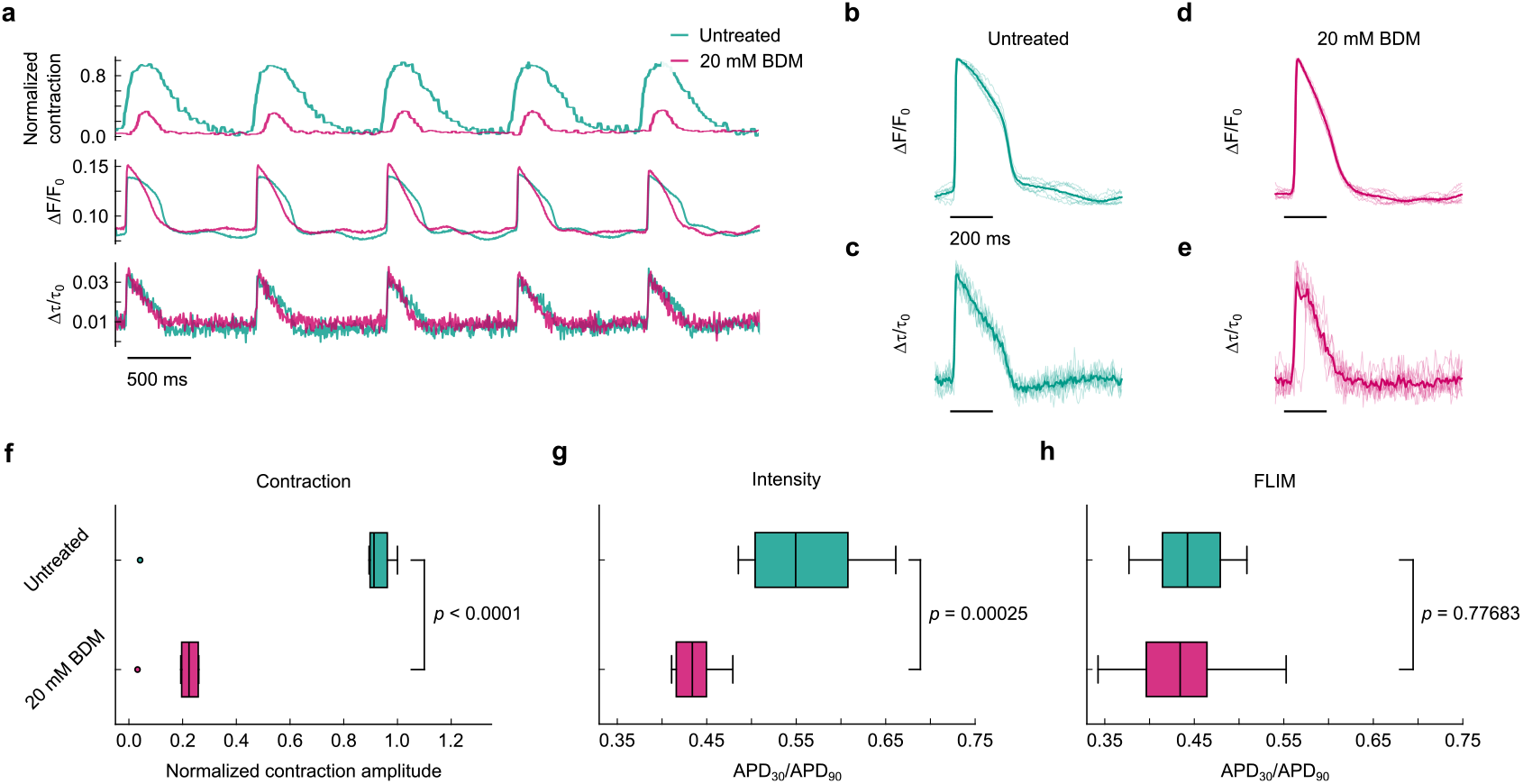
Comparison of fluorescence lifetime and intensity voltage measurements highlights motion-robust signals from lifetime imaging in human cardiac monolayers. (**a**) Excitation-contraction time traces in a hiPSC-CM monolayer paced at 1 Hz under natural contractile conditions (green) and after treatment with BDM (pink). Top to bottom: contraction waveforms, APs measured with intensity, and APs measured with fluorescence lifetime. (**b**-**e**) Time-averaged AP waveforms from intensity-(**b, d**) and lifetime-based (**c, e**) measurements. Individual traces are overlaid in a lighter shade to illustrate variability. (**f** ) Box plot analysis of the reduction in contraction amplitude induced by BDM treatment. (**g, h**) Box plot showing the change in *APD*_30_*/APD*_90_ before and after BDM treatment measured with fluorescence intensity (**g**) and fluorescence lifetime (**h**). For (**e**-**g**) the boxes show the IQR, the horizontal lines represent medians, and the whiskers extend to 1.5*×* the IQR.

AP time traces were recorded with FLOE from 204 × 204 µm FOVs (Fig. 2a middle and bottom) under baseline contractile conditions (green traces) and after treatment with 20 mM BDM (pink traces). Under natural contractile conditions, the Δ*F/F*_0_ recordings exhibit pronounced distortions in the shape and duration of the AP waveforms (Fig. 2a, middle, green), indicative of motion artifacts. This effect is highlighted in Fig. 2b where we show significant deviation between individual beats and the average waveform. This is substantially reduced after BDM treatment (Fig. 2a, middle, pink and d). In contrast, the Δ*τ/τ*_0_ measurements (Fig. 2a, bottom) show nearly identical AP waveforms recorded before and after BDM treatment (Fig. 2c, e respectively). We note that intrinsic fluorescent signals (e.g., from NADH, FAD) can generate weak autofluorescence signals, but their reported lifetimes are typically in the range of 300–600 ps [39], which is far shorter than the ns lifetimes we consistently measure. Combined with the fact that these endogenous signals are too dim to support reliable lifetime extraction under our imaging conditions, we are confident that the lifetime contrast reported here is dominated by voltage-sensitive responses rather than contamination from native fluorophores, with the same principle applying to the calcium signals discussed in later sections.

When looking at the *APD*_30_*/APD*_90_ ratio for individual beats, intensity-based measurements exhibited a highly significant shift after BDM treatment (*p <* 0.0005), demonstrating that contractile motion induces blur that obscures the true electrophysiological parameters when using traditional measurement techniques. In contrast, FLIM measurements in untreated samples (actively contracting) produced APD ratios statistically indistinguishable from those taken after motion inhibition (*p* = 0.78), showing that FLIM measurements accurately reflect AP activity in contracting monolayers. Moreover, the interquartile ranges (IQRs) for the untreated and BDM-treated FLIM datasets were comparable (0.68 and 0.65, respectively), underscoring FLIM’s intrinsic resilience to motion artifacts and its ability to capture physiologically relevant AP dynamics in contracting CMs with similar accuracy to that achievable after pharmacological inhibition. Additionally, as FLOE is expected to produce motion-insensitive waveform morphologies, post-treatment intensity and lifetime traces should exhibit strong agreement, as both should in principle be devoid of motion artifacts, as shown in Fig. 3d, e.

**FIG. 3.**
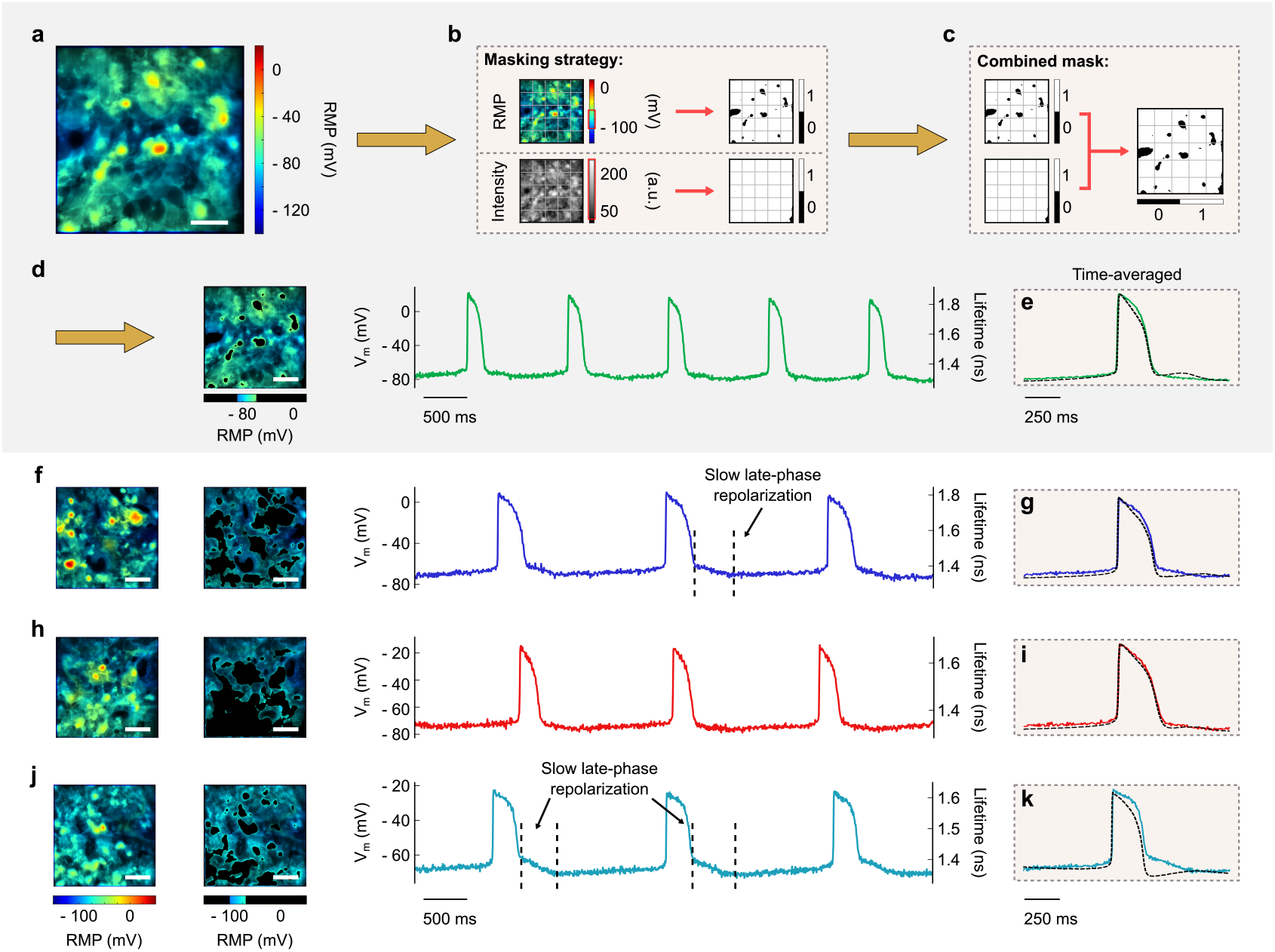
Calibrated mapping of voltage dynamics in intact human cardiac monolayers. (**a**) Spatially resolved map of the RMP during diastole. (**b**) Two-step masking strategy to remove regions with abnormal RMP (top) and low signal (bottom), ensuring that only cells within the physiological range are analyzed. (**c**) Combined mask used for all voltage-based analysis formed by combining the intensity and RMP masks. (**d**) Masked RMP map and voltage-based analysis showing spatially resolved RMP map and corresponding AP time trace. (**e**) Temporally averaged waveforms from (**d**) recorded with fluorescence lifetime (colored) and intensity (black dashed). (**f** -**k**) Equivalent analyses for 3 additional regions from two distinct monolayers: (**f, h**) correspond to one monolayer, while (**a, j**) correspond to another. White scale bar in all images, 25 µm.

Taken together, the reduction in variation in waveform morphology and *APD*_30_*/APD*_90_ ratios illustrates the inherent resilience of FLOE to motion artifacts.

### Quantitative analysis of regional action potential dynamics

To demonstrate the ability of FLOE to record absolute membrane potential values in hiPSC-CMs, we leveraged the linear dependence of the fluorescence lifetime of FluoVolt on membrane potential values [27]. We verified this with a multipoint calibration using valinomycin-treated H9C2 cells exposed to sequentially increasing potassium ion concentrations to establish a range of membrane potential values between −80 and 0 mV [40]. This calibration establishes a linear reference framework that closely matches previously reported values [27] and allows interpolation of V_m_ in hiPSC-CM monolayers for high-resolution electrophysiological analyses (see Methods and Supplementary Fig. 2 for details).

We imaged FluoVolt-stained hiPSC-CM monolayers and interpolated their V_m_ from the calibration curve. An example image of the resting membrane potential (RMP) is shown in Fig. 3a. To ensure that only physiologically relevant regions were included in our analyses, we applied a two-step masking strategy (shown pictorially in Fig. 3b, c). First, an intensity-based mask removed areas without cells or with insufficient signal (Fig. 3b bottom). Second, a mask based on RMP, retained cells with near normal RMP values ( −60 to−100 mV) during diastole (Fig. 3b top). Together, these masks produced maps that more accurately reflect functional heterogeneity within the monolayer (Fig. 3c, d). The resulting masked image was then used to analyze dynamic AP activity across the field (Fig. 3d). When overlapping the lifetime traces with the intensity traces (Fig. 3e), we see motion artifacts in the intensity trace (black) but not in the lifetime trace (green).

With the calibration in place, we then mapped AP dynamics across multiple spontaneously beating hiPSC-CM monolayers and observed significant spatial heterogeneity in waveform morphology (Fig. 3d-k). Regional differences in depolarization and repolarization kinetics were evident, highlighting functional variability within the monolayers. It is worth noting that some images (particularly Fig. 3f) show non-physical RMP values. In reality, all cells with compromised membranes should have a membrane potential of approximately 0 mV, but more positive values are seen as a result of saturation of the gated images. Some time traces (e.g., Fig. 3h, j) also exhibited lower than expected amplitudes, which may reflect contributions from adjacent uncoupled non-excitable cells that remain polarized and thus reduce the apparent average signal. A quantitative analysis of the time-averaged waveforms shown in Fig. 3e, g, i, k is given in Supplementary Fig. 3.

We also found that the signal-to-noise ratio (SNR) of the averaged lifetime waveforms (Fig. 3e, g, i, k colored) are comparable with that of the individual traces in Figs. 3d, f, h, j – underscoring our system’s robustness in capturing rapid variations from beat to beat with minimal noise interference.

### Distinct late repolarization dynamics

The level of fidelity in temporal resolution given by FLOE not only ensures accurate detection of dynamic changes in AP activity, it can also provide additional physiological information. The commercial hiPSC-CMs used in this study, and generally elsewhere, have an immature cardiac phenotype that differs from isolated adult ventricular CMs [41], including the ability to beat spontaneously – a feature that is absent in adult ventricular CMs [42–44]. The AP waveform shape and time course were previously reported using both optical dyes and glass micro-electrodes. However, in some instances here, the AP waveform contained a final late phase of repolarization with a considerably slower *dV/dt* (annotated examples in Fig. 3f, j). For the time-averaged AP trace shown in Fig. 3k, this slower phase of repolarization exhibits a gradient of -9 mV/ms. This feature has, to our knowledge, not been observed elsewhere. This is likely due to the fact that the intensity-based measurements normally used are unable to capture these late-stage dynamics because they are obscured by residual motion artifacts. We show this suppression in Fig. 3g, k by plotting the time-averaged FLIM and intensity readings from the sample in Fig. 3f, j, where this distinct slow repolarization is captured in lifetime measurement but not intensity.

### Spatiotemporal dynamics of calcium transients in hiPSC-CM monolayers

To show FLOE’s ability to measure electrophysiological changes over a large FOV, we recorded Ca^2+^ transients in spontaneously firing hiPSC-CM monolayer samples stained with Cal-520, AM.

In Fig. 4a, b, we show example intensity and lifetime plots, respectively. When expanding to a large FOV (1.6 × 1.6 mm) the illumination profile causes variations in the intensity levels across the field as shown in Fig. 4c. In contrast, FLOE provides a uniform life-time profile across the FOV, as would be expected from a intact monolayer (Fig. 4d). It is worth noting that like the AP recordings, exceedingly low intensity values cause inaccurate lifetime estimations, as shown in Fig. 4e.

**FIG. 4.**
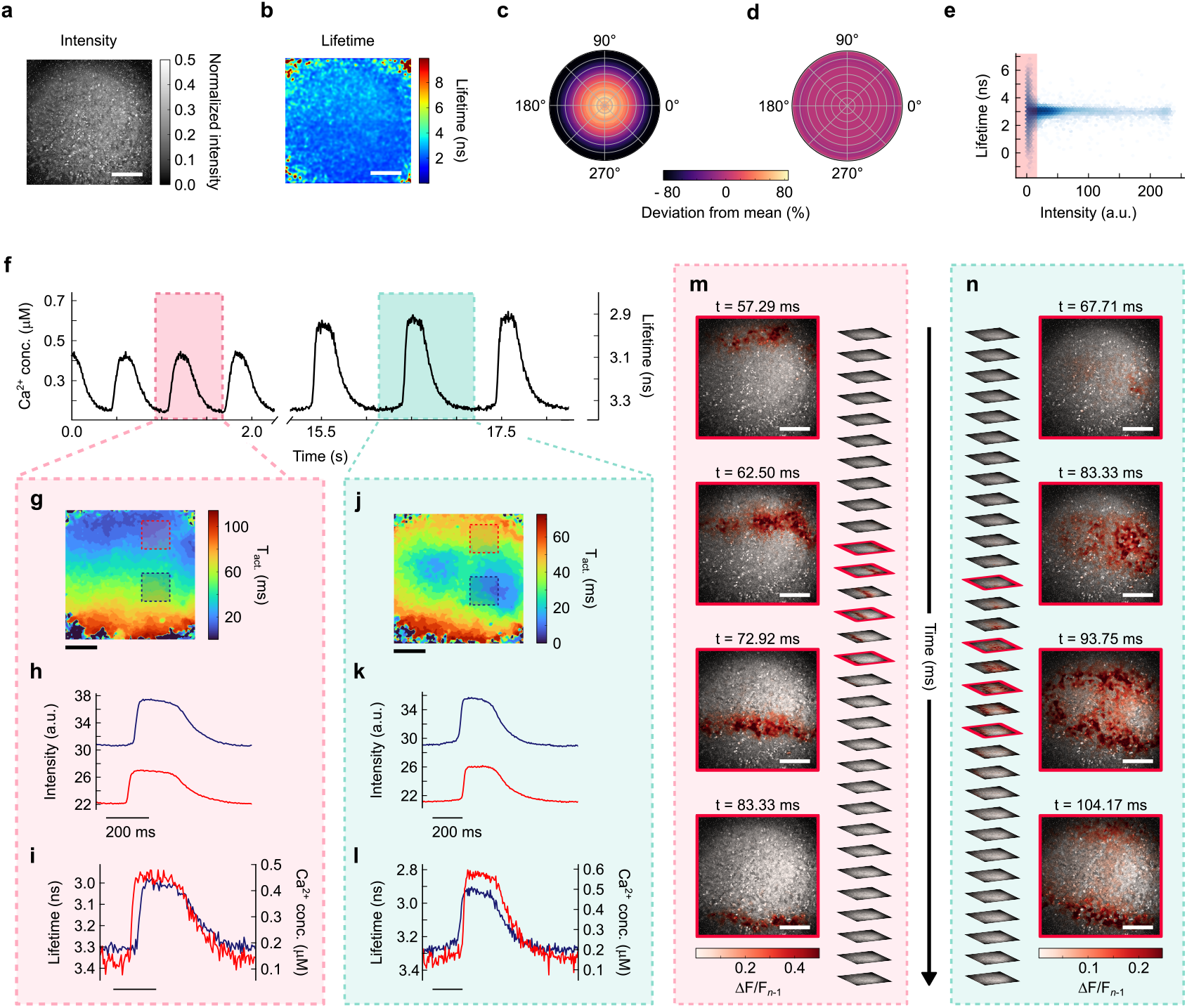
Spatiotemporal mapping of calcium wave propagation in hiPSC-CM monolayers by widefield fluorescence lifetime imaging. (**a, b**) Representative fluorescence intensity (**a**) and lifetime (**b**) images of a hiPSC-CM monolayer over a 1.6 × 1.6 mm FOV. White scale bar, 400 µm. (**c**,**d**) Polar plots showing percentage deviation from mean intensity (**c**) and lifetime (**d**) across the images in (**a, b**) respectively. (**e**) Plot of lifetime versus intensity, with the red region indicating the same low-intensity pixels as masked in voltage analyses to ensure stable lifetimes. (**f** ) Representative calcium transient time trace. (**g**-**i**) Activation map of the first beat highlighted in pink in (**f** ) showing the time of calcium activation at each spatial region (T_act._). The activation wave propagates from top to bottom. (**h**), intensity traces from two regions of interest highlighted in (**g**) and (**i**), the corresponding lifetime traces. (**j**-**l**) Same as (**g**-**i**) for the second beat highlighted in green in (**f** ). The activation wave now propagates from central regions, outwards. Black scale bar in (**g, j**), 400 µm. (**m**,**n**) Frame-by-frame visualization of calcium wave propagation for the beats in (**g, j**), with highlighted frames outlined in red. Colors of surrounding boxes match traces in (**f** ). Images depict instantaneous fractional intensity changes (Δ*F/F*_*n*−1_). White scale bar, 400 µm.

Similarly to the calibration of the AP values, we can also provide a direct calibration of the fluorescence lifetime values to intracellular ion concentrations, allowing quantitative mapping of Ca^2+^ transients with high spatiotemporal precision. To perform this calibration, we imaged mixtures of Cal-520, potassium salt in solution with varying concentrations of Ca^2+^ ions to produce a calibration curve and interpolate ionic concentrations in hiPSC-CM monolayers during spontaneous activity (see Methods for details). The lifetime values estimated with our RLD approach show strong agreement with previously reported measurements for fluorescein-based, BAPTA-derived Ca^2+^ indicator dyes [45], measured in the corresponding green spectral range, consistent with the expected photophysical behavior of Cal-520. A representative time trace illustrating the dynamic intracellular Ca^2+^ concentrations during spontaneous activity is shown in Fig. 4f.

Quantitative analysis of individual Ca^2+^ waveforms and related parameters is typically required to evaluate cell health from the kinetics of calcium handling. We applied this analysis to a representative transient located halfway through the recording shown in Fig. 4f. The results of this analysis are shown in Table II in the Supplementary Material.

When isolating two individual transients from the recording in Fig. 4f and examining their activation maps (Fig. 4g, j), we see dynamic changes in the position of the pacer cells and their wave propagation. More specifically, Fig. 4g shows a single wavefront propagating from the top of the FOV down, whereas Fig. 4j depicts two concentric rings originating from distinct points and propagating radially outward. Here, T_act._ is defined as the time between the baseline and 50% the upstroke. When looking at two distinct regions with different activation times, we see that the unprocessed intensity traces (Fig. 4h, k) show different baseline intensity levels and varying amplitudes due to non-uniform illumination, as the trace from the red region shows a consistently lower baseline intensity than the blue region. This is not the case when examining the lifetime traces (Fig. 4i, l), which show nearly identical amplitudes and baselines agnostic of the sample location. Here, activation maps are generated from intensity recordings to maximize SNR. Although lifetime data produce similar activation profiles (Supplementary Fig. 4), we emphasize that lifetime and intensity measurements offer complementary strengths and a thoughtful combination of both provides the most robust analysis. While lifetime imaging offers higher fidelity for resolving individual transients, intensity-based measurements remain the preferred approach for constructing activation maps. We argue that lifetime will not replace intensity as the universal standard; rather, the most powerful insights will come from a deliberate combination of both modalities.

By plotting the instantaneous change in fluorescence lifetime or intensity between each subsequent frame instead of comparing to a resting baseline (notated here as Δ*τ/τ*_*n*−1_ and Δ*F/F*_*n*−1_, respectively), we can also visualize the position of the Ca^2+^ wavefront during periods of spontaneous cardiac activity. Example spatially resolved recordings of the transients highlighted in Fig. 4f are shown in Fig. 4m, n. Here, we present intensity recordings obtained using FLOE’s dual-modality data capture, which maximizes the SNR. Similarly, spatially resolved wavefront videos of AP propagation can be generated for FluoVolt-stained samples. These recordings complement calcium imaging by capturing the dynamics of electrical activity throughout the monolayer, providing a comprehensive view of the underlying excitation processes (see Supplementary Material Fig. 5 for additional results).

By capturing subtle beat-to-beat variations in Ca^2+^ dynamics, we demonstrate FLOE’s ability to perform precise quantification of the rapid Ca^2+^ wave, providing a robust metric to assess excitation-contraction coupling.

## DISCUSSION

FLOE delivers rapid, noninvasive voltage and calcium ion mapping with high spatial resolution while inherently rejecting motion artifacts from native contractile behavior. This high-fidelity approach thus circumvents the artifacts present in traditional intensity-based imaging, offering a powerful tool for spatiotemporal analysis of hiPSC-CM electrophysiology. In particular, the study shows that recording simultaneously both intensity and lifetime signals allows the exploitation of their relative strengths when analyzing and interpreting signals from voltage sensitive and calcium sensitive fluorescent probes.

As an example, we report an unexpected feature in the late-phase AP repolarization that exhibits a slower negative gradient. This appears to vary in prevalence across the sample and has most likely been previously overlooked due to motion artifacts in standard intensity measurements. We can only speculate on the underlying mechanism, but one possible cause may be related to intracellular calcium transient as the decay phase of this transient extends beyond the repolarizing phase 3. The decay of the intracellular calcium transient is mediated by the activity of the electrogenic sodium-calcium exchanger (NCX) [18, 46]. In forward mode, this generates an inward current that would explain the late-phase potential [47]. Examples of this waveform are present in two of the four recordings shown. The reason why this characteristic is seen in some, but not all, recordings is unclear, but cell-to-cell differences in electrophysiology and intracellular calcium transients are a feature of this cell type and this may represent the range of electrophysiological phenotypes present [48, 49].

Importantly, quantitative analysis (Fig. 2e) reveals that FLOE measurements in contracting CMs yield statistically comparable *APD*_30_*/APD*_90_ ratios (*p* = 0.77683) between contracting and non-contracting preparations. The difference between AP waveforms before and after BDM is due to two effects: (i) inhibition of contraction and loss of movement artifact, and (ii) direct action of BDM on ion channels [50] that changes the shape and duration of the action potential. Intensity measurements cannot distinguish these effects. Comparison of AP intensity and lifetime waveforms in a contracting monolayer reveals differences in waveform shape due to motion artifacts in the intensity trace. Comparing intensity and lifetime measurements after BDM indicates a closer correspondence of the time-courses since, in both cases, the change in signal is entirely due to voltage. These findings emphasize that FLOE can reliably capture physiologically relevant AP dynamics in actively contracting cells without the need for pharmacological motion suppression.

Additionally, we demonstrate that FLOE provides sufficient SNR to allow single-beat analysis without temporal averaging, providing a more physiologically relevant assessment of cardiac excitability [51, 52].

Furthermore, resolving single-beat characteristics establishes a foundation for studying arrhythmic events and cellular responses to pharmacological agents. The observed heterogeneity in the AP waveforms between individual beats and sample regions suggests that regional differences in the ion channel activity may contribute to functional disparities within the monolayer. Future work integrating pharmacological perturbations could further explore the mechanisms underlying this variability [53, 54].

The versatility of FLOE extends beyond cardiovascular research. For example, in neuronal electrophysiology, where precise voltage mapping is crucial to understanding synaptic transmission and network behavior, our method could provide critical insights without the need for invasive electrodes [34, 55–5 . Similarly, this setup could be adapted to study other tissues where rapid fluctuations in environmental factors are indicative of cellular function, e.g., to monitor dynamic processes using tension probes in mechanobiology [58]. The ease of integrating this approach into existing microscope platforms further enhances its appeal, as it avoids the complexity and cost associated with other fast FLIM setups [33, 34].

Despite these advantages, we acknowledge limitations of our imaging system in its current configuration which should be addressed in future iterations. All data presented were acquired at 192 fps, which is slower than current intensity-based techniques which have achieved frame rates exceeding 1,000 fps, although with varying levels of spatial resolution [19].

Future work will indeed focus on improving frame rates by leveraging lower spatial resolution but faster SPAD cameras in combination with high-resolution CMOS cameras to computationally recover resolution [59] or by investigating the opportunity offered by SPAD cameras to operate at faster frame rates by reducing the bit-depth. Additionally, dual imaging of voltage and calcium dynamics would be made possible by using a second SPAD camera. This would require independent lifetime measurements based on spectral separation of the dyes – possible with the use of FluoVolt and Cal590 [60]. This approach would allow simultaneous recording of all three aspects of excitation-contraction coupling – currently not possible with existing technologies.

In conclusion, we believe that FLOE provides a significant advantage over alternative methods for studying cardiac electrophysiology and other conductive tissues without compromising spatial resolution or relying on motion uncouplers.

## METHODS

### hiPSC-CM culture and preparation

Commercially sourced hiPSC-CMs were cultured under controlled conditions (5% CO_2_, 37^°^C) following the optimized protocols of the manufacturer (Celogics). Cells were plated onto fibronectin-coated (Sigma-Aldrich, F1411) 35 mm glass-bottom dishes (CellVis, D35-14-1.5-N) using plating medium. The following day, the medium was replaced with maintenance medium, with subsequent media changes performed every 48 hours. For functional imaging, cells were stained with either 1 µM Cal-520, AM (AAT Bioquest, 21130) with 0.02% Pluronic F-127 (Biotum, 59000) to assess intracellular calcium dynamics, or 1:1000 FluoVolt (ThermoFisher Scientific, F10488) and 1:100 powerload for AP recordings. The dyes were loaded for 30 minutes, followed by a single wash with maintenance medium. Cells were then incubated under standard culture conditions until imaging. All recordings were conducted in a controlled environment using an on-stage incubator to maintain physiological conditions throughout data acquisition.

### Microscope setup and imaging conditions

The fluorescence imaging system was based on an inverted fluorescence microscope design with a split output port to allow simultaneous imaging on two cameras: a large-format SPAD array for FLIM and a scientific CMOS for high-resolution intensity imaging. Excitation was provided by a diode laser operating at 80 MHz, with a peak wavelength of 472 ± 7 nm, a typical pulse width of 65 ps and a maximum pulse width of 90 ps (HORIBA Scientific, DeltaDiode-L, 470 nm). The light was delivered through a 150 µm^2^ square-core multimode fiber (Thorlabs, M101L02) to ensure uniform illumination and minimize intensity variations from a Gaussian profile. To reduce speckle patterns in the illumination, the fiber was vibrated during experiments with a small motor. This illumination scheme was originally proposed in [61]. The excitation beam was focused onto the back aperture of an objective using a 100-mm achromat (Thorlabs, AC254-100-A-ML). The objective was mounted on a piezo-controlled stage to enable precise focus adjustments. Emission signals were collected and separated from excitation light using a FITC dichroic mirror (Thorlabs, MD499) (reflection band: 470-490 nm, emission band: 508–675 nm), followed by additional spectral filtering with a 525/50 nm bandpass filter (Chroma Technology Corp., ET525/50 M). The cleaned emission signal was then split with a 70:30 beamsplitter and imaged, respectively, onto the SwissSPAD3 [37] and a 10-megapixel scientific CMOS (Teledyne Photometrics, Kinetix). A red LED was also mounted above the objective to provide brightfield imaging capabilities (Thorlabs, M780LP1).

The magnification of the microscope was controlled by alternating objective and tube lenses to suit specific imaging requirements. A 40× water immersion objective (Nikon, 40× /1.15 NA) was used in combination with a 200 mm tube lens (Thorlabs, TTL200-A) for all 40× imaging. Air immersion 10× (Nikon, 10×/0.50 NA) and 20× objectives (Nikon, 20×/0.75 NA) were used with a 100 mm tube (Thorlabs, AC508-100-A) lens to demagnify the system by 2× . This provided wider FOVs whilst improving light collection by maintaining relatively high NAs with higher magnification objectives.

To maintain physiological conditions, a heated incubator (Okolabs, H301-K-FRAME) was mounted on the system above the objective to house the samples during imaging. The samples were exposed to 5% CO_2_ and maintained at 37^°^C for the duration of all experiments. The incubator was mounted on a motorized XY microscope stage (Thorlabs, MLS203-1) to allow fine control over sample position.

The SPAD array has an 8-bit frame rate of 192 fps with an exposure time of 10.24 µs per binary frame. 8-bit images were generated by collecting 255 binary frames which were summed on the FPGA before reading out. The CMOS camera was configured to acquire 12-bit images at 200 fps with a 5 ms frame exposure. To approximately match the field of view of the two sensors, the larger CMOS was cropped to a 1400 × 1400-pixel region of interest. The two sensors were run synchronously by triggering each CMOS frame with SPAD. All FLIM data were captured with the SPAD, with the CMOS used only for contractility measurements.

### Membrane potential calibration via fluorescence lifetime imaging

For calibration of fluorescence lifetime to membrane potential values, H9C2 cells were seeded in 35 mm glass bottom plates (Cellvis, D35 20 1.5H) precoated with fibronectin (Sigma-Aldrich, F1141) and maintained in standard growth medium until 70-80% confluence. These H9C2 cells were obtained as a gift from Prof. George Baillie (Cardiovascular & Metabolic Health/University of Glasgow). For membrane potential measurements, cells were washed and incubated in Tyrode’s solution (92.3 mM NaCl, 20 mM NaHCO_3_, 1 mM Na_2_HPO_4_, 1 mM MgSO_4_, 5 mM KCl, 20 mM sodium acetate, 25 mM glucose, 5 mM HEPES, 1.8 mM CaCl_2_; pH 7.4). During imaging, samples were maintained at 37^°^C under 5% CO_2_ in an on-stage incubator. Baseline fluorescence lifetime recordings were acquired on the FLIM system. The cells were then treated with 1 µM valinomycin (Sigma-Aldrich, 94675) for 10 minutes to hyperpolarize the membrane and a second recording was obtained. Subsequently, extracellular KCl was increased stepwise—from 2.7 mM to final concentrations of 8, 12, 20, 36 and 140 mM—to impose calculated potentials of –72, –65, –47, –32, and 0 mV, respectively. At each K^+^ concentration, cells were allowed to equilibrate for 2 minutes prior to imaging. Data were analyzed as previously described [40].

### Calibration of fluorescence lifetime to intracellular calcium concentrations

For calibration of fluorescence lifetime measurements with defined intracellular Ca^2+^ levels, calibration solutions were prepared using the calcium-sensitive dye Cal-520, AM (potassium salt; AAT Bioquest, 21140) in buffers with precisely controlled free Ca^2+^ concentrations.

Calibration of calcium concentrations was achieved using Ca-EGTA and EGTA buffers. Briefly, 100 mM Ca-EGTA and 100 mM EGTA were prepared separately by mixing each with 500 mM HEPES (pH 7.0 at 21^°^C) and ultrapure water in a 1:1:8 ratio. 1 µM Cal-520 potassium salt (AAT Bioquest, 21140) was added to each solution. Ca-EGTA and EGTA were then mixed as follows to obtain a range of buffer solutions with known calcium concentrations: Ca-EGTA + 10 µM CaCl (25 µM Ca^2+^), Ca-EGTA only (15 µM), 10:1 (4 µM), 3:1 (820 nM), 1:1 (400 nM), 1:3 (82 nM), 1:10 (40 nM) and EGTA only (0.2 nM).

For imaging, 200 µL of each solution was dispensed onto a 96-well glass bottom plate and fluorescence lifetime measurements were acquired using the FLIM system. The resulting calibration curve (Supplementary Fig. 6) enabled precise interpolation of intracellular Ca^2+^.

### Contractility assessment

The contractile function was evaluated using high-speed bright-field imaging. The images were acquired at 200 fps with a 40× objective to ensure high temporal resolution of dynamic cellular contractions. Graphite electrodes were used to field stimulate the hiPSC-CMs at a fixed rate of 1 Hz and to synchronize beating across the culture. In a subset of experiments, contraction was uncoupled by administering 20 mM BDM.

The acquired image stacks were analyzed using the MUSCLEMOTION algorithm [38] to quantitatively assess contractile parameters.

### Rapid lifetime determination algorithm

To generate spatially resolved fluorescence lifetime maps, we employed RLD using the dual-gated capabilities of SwissSPAD3 [37]. This sensor builds images by capturing binary frames of photon detection events and summing consecutive frames to produce images with a higher bit depth. For all results presented, the SPAD was operating in 8-bit mode resulting in an acquisition rate of 192 fps. For each exposure, SwissSPAD3 returns both a time-integrated intensity image (*INT* ) and a gated image (*G*2). An antigated image (*G*1) is also computed as the difference between these, *G*1 = *INT* − *G*2. It is worth noting that, while an ideal SPAD gate is often modeled as a rectangular (boxcar) function, in practice the gate and antigate exhibit finite rise and fall times and a slightly rounded plateau. For all results presented in this study, a gate width approximately equal to the expected lifetime was used for *G*2. In the monolayer samples analyzed here, FluoVolt typically exhibits a fluorescence lifetime of *τ* ∼ 1.35 ns at rest and *τ* ∼ 1.7 ns during peak AP activity, while Cal-520, AM exhibits a fluorescence lifetime of *τ* ∼ 3.3 ns during rest periods and *τ* ∼ 2.9 ns at maximum Ca^2+^ activity. To build a lookup table relating the gate-to-antigate ratio and fluorescence lifetime, we simulate normalized fluorescence decay functions of the form

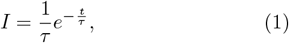

with varying lifetimes, *τ*, for *t* ≥ 0 and *τ >* 0 and compute the dot product between each decay and the SPAD’s gate profile. An example lookup table is shown in Supplementary Fig. 7c.

### Data processing and temporal cleaning

Before any processing is performed, all raw images undergo pile-up correction to account for undetected photons arising from the binary acquisition of the sensor. The total detected counts *C*_*D*_ can be estimated from the measured counts *C*_*M*_ following

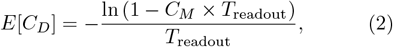

where *T*_readout_ = 6.4 µs for the SwissSPAD [62].

We perform background subtraction on all corrected data (gate, antigate and intensity) on a pixel-wise basis using dark frames averaged over a 40 s acquisition. For data with particularly low light levels, we also increased the SNR by spatially blurring each frame. To maintain image dimensions and avoid loss of fine structures, this is done by convolving each image with a two-dimensional Gaussian kernel of the form

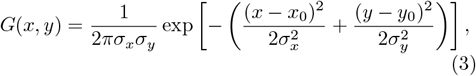

where *σ*_*x*_ and *σ*_*y*_ represent the standard deviations in *x* and *y* respectively. For all data presented, we use a 20 × 20 kernel size with *σ*_*x*_ = *σ*_*y*_ = 4.

Following fluorescence lifetime computation, we process the FLIM data further to suppress spatial noise caused by fluctuations in photon levels in the temporal domain. This method of temporal cleaning, shown in [63], fits an analytical expression to the three-dimensional data cube along the temporal dimension for each spatial co-ordinate pixel and thresholds out areas with no activity. This temporal cleaning was applied only to the data visualized as Δ*τ/τ*_*n*−1_ (Fig. 4m, n and Supplementary Fig. 4a, 5a). For all cardiac data shown, a skew-normal distribution

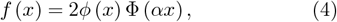

with shape parameter *α* was used to approximate the calcium and AP waveforms. In Eq. 4, *ϕ* (*x*) represents the standard normal probability density function

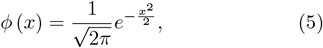

and Φ (*x*) is the cumulative distribution function

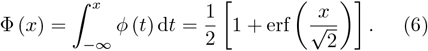

The fitting was restricted to the amplitude, *t*_0_ and shape parameters for each spontaneous firing event, and dormant pixels were identified and zeroed using these free parameters (see Supplementary Fig. 9 for additional details). In this way, the timing information contained in each pixel allows for enhancement to the spatial quality of our data.

## ACKNOWLEDGEMENTS

We thank I. Starshynov for his assistance with porting the camera control code used in this study, S. B. Forbes for providing the immunofluorescence images, and E. McGhee and F. L. Burton for insightful discussions. The authors acknowledge funding from the Engineering and Physical Sciences Research Council and UKRI (UK, grant nos. EP/Z533166/1, EP/T00097X/1, EP/Y029097/1 and EP/V051148/1). V.Z. acknowledges funding from the Research Council of Lithuania (project no. S-MIP-24-91). E.H. and G.L.S. received support from the British Heart Foundation (special project grant no. SP/F/23/150059), and C.M. and G.L.S. from the British Heart Foundation (grant no. NH/F/21/70005). J.Z., G.G.T. and E.C. acknowledge support from the U.S. Department of Energy (grant no. DE-SC0023184). D.F. is supported by the Royal Academy of Engineering through the Chairs in Emerging Technologies program.

## AUTHOR CONTRIBUTIONS

E.M.: conceptualization, methodology, investigation, formal analysis, visualization, writing - original draft. V.Z.: conceptualization, methodology, supervision, writing - review & editing. E.H.: methodology, investigation, resources, writing - original draft. G.A.: investigation. J.Z.: software, writing - review & editing. G.G.T.: software, writing - review & editing. C.B.: resources, writing - review & editing E.C.: resources, writing - review & editing. G.L.S.: conceptualization, methodology, supervision, writing - review & editing. C.M.: conceptualization, methodology, supervision, writing - review & editing. D.F.: conceptualization, methodology, supervision, funding acquisition, writing - review & editing.

## COMPETING INTERESTS

G.L.S. is co-founder, director, board member, and Chief Scientific Officer of Clyde Biosciences Ltd. (UK). The remaining authors declare no competing interests.

## DATA AVAILABILITY

All data supporting the findings of this study will be made available upon publication.

