## Supplemental information for "Fluorescence-lifetime optical electrophysiology in contracting cardiomyocytes"

### Supplementary material: Fluorescence-lifetime optical electrophysiology in contracting cardiomyocytes

Euan Millar[1], Vytautas Zickus[1,2], Eline Huethorst[3], Giedrė Astrauskaitė[1], Jiuxuan Zhao[4], Gregor G. Taylor[4], Claudio Bruschini[4], Edoardo Charbon[4], Godfrey L. Smith[3], Caroline Müllenbroich[1,*], Daniele Faccio[1,*]
[1]*School of Physics and Astronomy, University of Glasgow, Glasgow G12 8QQ, UK*
[2]*Department of Laser Technologies, Center for Physical Sciences and Technology, LT-10257, Vilnius, Lithuania*
[3]*School of Cardiovascular and Metabolic Health, University of Glasgow, Glasgow G12 8QQ, UK*
[4]*Advanced Quantum Architecture Laboratory (AQUA), École Polytechnique Fédérale de Lausanne (EPFL), Neuchâtel 2002, Switzerland*

#### DATA VISUALIZATION

All fluorescence lifetime images shown (Fig. 3e-h in the main paper) are visualized by weighting (overlaying) the raw FLIM data with the corresponding intensity image (also captured by the SPAD). This is done by first mapping the lifetime values to RGB colors using a chosen colormap, producing a false-color image that represents the spatial distribution of fluorescence lifetimes. Separately, the associated intensity image is first processed by filling in dead pixels in the SPAD. These were identified by comparing the median-filtered versions of the image (with kernel sizes 3 and 11), and replaced by using the lower scale ($3 \times 3$) median values to fill regions with abnormally high variation. Then, contrast limited adaptive histogram equalization (CLAHE) is applied. This enhances local contrast throughout the field of view (FOV) while avoiding overamplification of noise or saturation of bright regions. Finally, each RGB channel of the lifetime image is scaled by the enhanced intensity image, preserving the original color (lifetime) information but adjusting the brightness according to local intensity. This combined visualization improves the interpretability of spatial lifetime features compared to the raw FLIM images.

The intensity images shown with no FLIM overlay (Fig. 2c, d in the main paper) were also processed using CLAHE to enhance contrast for visualization purposes.

To compare fluorescence lifetime changes under experimental conditions, we first employ a classical normalization approach relative to a baseline frame, defined as

$$\Delta\tau/\tau_0 = \frac{\tau_n - \tau_0}{\tau_0}, \tag{1}$$

where $\tau_n$ denotes the fluorescence lifetime in frame $n$, and $\tau_{\text{baseline}}$ is the mean pre-stimulus or reference lifetime. This global $\Delta\tau/\tau_0$ metric is widely used in electrophysiology studies to quantify absolute changes over time and is effective in assessing action potential and calcium waveforms, as shown in Supplementary Fig. 5b.

To isolate and visualize propagating features such as lifetime wavefronts, we additionally compute the normalized frame-to-frame gradient

$$\Delta\tau/\tau_{n-1} = \frac{\tau_n - \tau_{n-1}}{\tau_{n-1}}. \tag{2}$$

This local temporal derivative emphasizes dynamic transitions between consecutive frames, enhancing the visibility of moving wavefronts, as shown in Supplementary Fig. 5a and Fig. 4a of the main paper. This method is particularly useful for highlighting spatially confined transient changes that may not be apparent on the $\Delta\tau/\tau_0$ map.

#### STATISTICAL ANALYSIS OF PHARMACOLOGICAL CONTRACTION SUPPRESSION

In the following, we show the statistical analysis performed to quantify the reduction in contraction parameters after treatment of the cardiomyocyte monolayers with 20 mM 2,3-butanedione monoxime (BDM) corresponding to Fig. 2 of the main paper. These data were analyzed with the open source software MUSCLEMOTION [1]. The box plots in Supplementary Fig. 1a show a comparison between the four main physiological parameters before and after BDM treatment. The four parameters analyzed are contraction duration at 50% amplitude ($\text{CD}_{50}$), contraction duration at 90% amplitude ($\text{CD}_{90}$), rise time ($\text{T}_{\text{Rise}}$), and decay time ($\text{T}_{\text{Decay}}$). The corresponding values are annotated on Supplementary Fig. 1b, c for the sample before and after BDM treatment, respectively. The two averaged waveform plots shown are normalized with respect to control (contractile) conditions. Due to the fact that both data sets were taken with the same illumination conditions, we can also compare the amplitudes of the normalized plots in Supplementary Fig. 1b, c to determine that the BDM treatment reduced the contraction amplitude of the monolayer by 76.27%. For all analysis shown, temporal sample sizes of $n = 8$ beats were used for

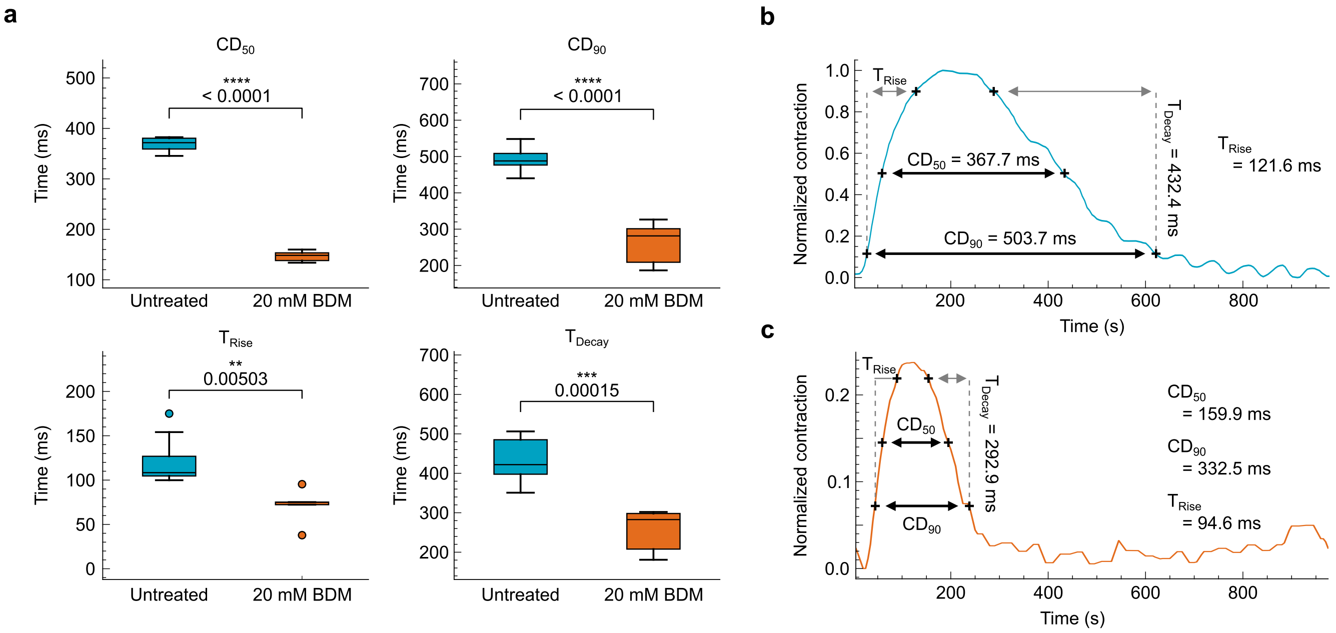


Supplementary Fig. 1. **Quantitative analysis of pharmacological contraction suppression.** (**a**) Quantitative comparison of contractile parameters—contraction duration at 50% amplitude ($CD_{50}$), contraction duration at 90% amplitude ($CD_{90}$), rise time ($T_{Rise}$), and decay time ($T_{Decay}$)—before and after BDM treatment. $n = 8$ for control and 5 for BDM-treated groups. The boxes show the IQR, the horizontal lines represent medians, and the whiskers extend to 1.5× the IQR. P-values are indicated directly on the plots, with significance levels denoted as follows: $P < 0.01$ (**), $P < 0.001$ (***), and $P < 0.0001$ (****). (**b**) Average contraction waveform of a untreated sample, annotated with the parameters analyzed in (**a**). (**c**) Average contraction waveform of a sample after motion suppression with 20 mM BDM, annotated with the parameters analyzed in (**a**).

control and 5 beats for the BDM-treated group, and both data were averaged over a 204 × 204 µm FOV, completely populated with cells (see Fig. 2b, c of the main paper).

The p- and t-values for the four parameters analyzed are summarized in Table I .

| Parameter | t-value | p-value |
|---|---|---|
| $CD_{50}$ | 28.66 | $1.09 \times 10^{-11}$ |
| $CD_{90}$ | 8.82 | $2.55 \times 10^{-6}$ |
| $T_{Rise}$ | 3.49 | $5.02 \times 10^{-3}$ |
| $T_{Decay}$ | 5.62 | $1.54 \times 10^{-4}$ |

TABLE I. t-values and p-values for all contraction parameters compared before and after BDM treatment.

#### CALIBRATION OF FLUORESCENCE LIFETIME TO MEMBRANE POTENTIAL

In this section, we provide additional details on the calibration procedure used to translate fluorescence lifetime values into membrane potential ($V_m$) readings, following protocols presented in [2]. Supplementary Fig. 2 shows a scatter plot in which each point represents the mean fluorescence lifetime measured in an H9C2 cell at a given $V_m$ (cells were in a confluent monolayer so the numbers varied between imaging regions but $n > 10$ cells were analyzed per polarization condition). The relationship between $V_m$ and the fluorescence lifetime was fitted with a linear regression which serves as the reference framework to interpolate $V_m$ in hiPSC-CM monolayers.

#### QUANTITATIVE ANALYSIS OF ACTION POTENTIAL WAVEFORMS

To quantify AP variations, we compared the averaged AP waveforms for multiple ROIs within two hiPSC-CM monolayers from the same plating. The table in Supplementary Fig. 3 reports the key parameters for the time-averaged waveforms from the traces shown in Fig.3 of the main paper, where early, middle and late repolarization are defined as the duration from APD 10-30, 40-60, and 70-90, respectively. These parameters are annotated on an example time-averaged trace in Fig. 3. As can be seen, the values for any given single AP waveform can differ

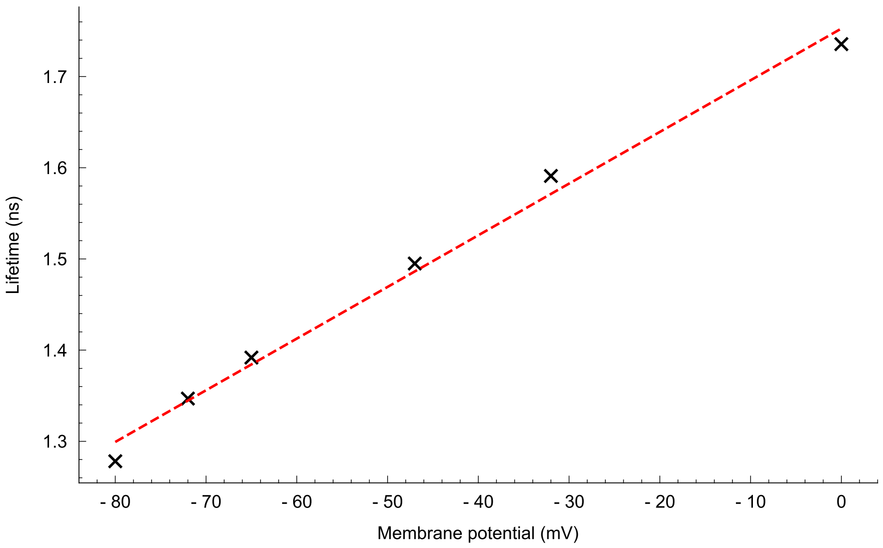


Supplementary Fig. 2. **Multipoint calibration of fluorescence lifetime versus membrane potential in H9C2 cells.** Fluorescence lifetime of FluoVolt–loaded H9C2 cells measured under discrete valinomycin-clamped membrane potentials ranging from –80 to 0 mV. A linear regression (red dashed line) gives a calibration curve for interpolation of membrane potential readings from fluorescence lifetime recordings.

c

| | Monolayer 1 (ms) | | Monolayer 2 (ms) | |
|---|---|---|---|---|
| | ROI 1 | ROI 2 | ROI 1 | ROI 2 |
| $APD_{90}$ | 337.14 | 254.01 | 362.09 | 213.22 |
| $T_{Rise}$ | 17.09 | 18.48 | 14.75 | 14.80 |
| Early rep. | 111.68 | 85.51 | 89.69 | 58.79 |
| Medium rep. | 33.11 | 26.11 | 24.50 | 17.84 |
| Late rep. | 29.48 | 25.08 | 71.48 | 25.07 |

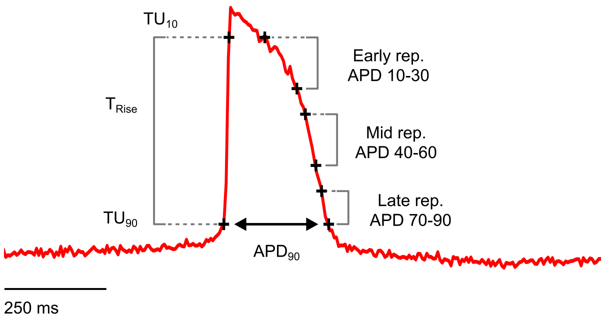


Supplementary Fig. 3. **Comparison of action potential parameters across multiple regions of two monolayers from a single plating.** Table of action potential waveform values quantitatively analyzed for the data shown in Fig. 3 of the main paper alongside a representative time-averaged waveform (Fig. 4h in the main paper) annotated with the parameters analyzed.

within the same sample. The data reported in the table were calculated from interpolated time traces to remove discretization from the camera frame rate.

#### IMAGING CALCIUM WAVE PROPAGATION WITH FLUORESCENCE LIFETIME

In Table II we show the quantitative analysis of a representative calcium transient from midway through the recording in Fig. 4f in the main paper.

| Parameter | Value (ms) |
|---|---|
| $CaT_{90}$ | 494.79 |
| $CaT_{50}$ | 296.88 |
| $T_{Rise}$ | 83.33 |
| $T_{Decay}$ | 286.46 |

TABLE II. Calcium transient parameters measured in a representative beat located halfway through the recording in Fig.4f of the main paper.

Furthermore, to directly compare the fluorescence lifetime analysis with conventional intensity-based readouts, we examined the propagation of calcium waves in the cardiomyocyte monolayers using the fluorescence lifetime signal. Although the main text (Fig. 4) shows the frame-by-frame dynamics of $\Delta F/F$ and the corresponding activation map extracted from the intensity data, here we present an analogous analysis performed on the lifetime

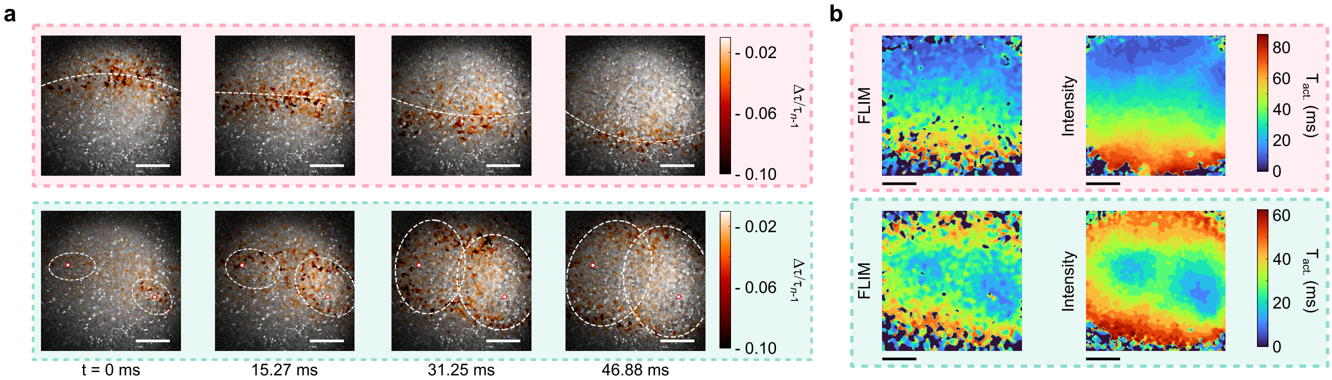


Supplementary Fig. 4. **Calcium wave propagation analyzed from fluorescence lifetime signals.** (**a**) Snapshots of $\Delta\tau/\tau_{n-1}$ frames corresponding to the time trace in Fig. 4 of the main paper. Colored boxes (pink, green) highlight the same beats marked in Fig. 4. (**b**) Activation maps showing the time of calcium activation ($T_{act.}$) extracted from fluorescence lifetime (left) and intensity (right). Both approaches reveal consistent propagation patterns, though with reduced signal-to-noise in the lifetime data.

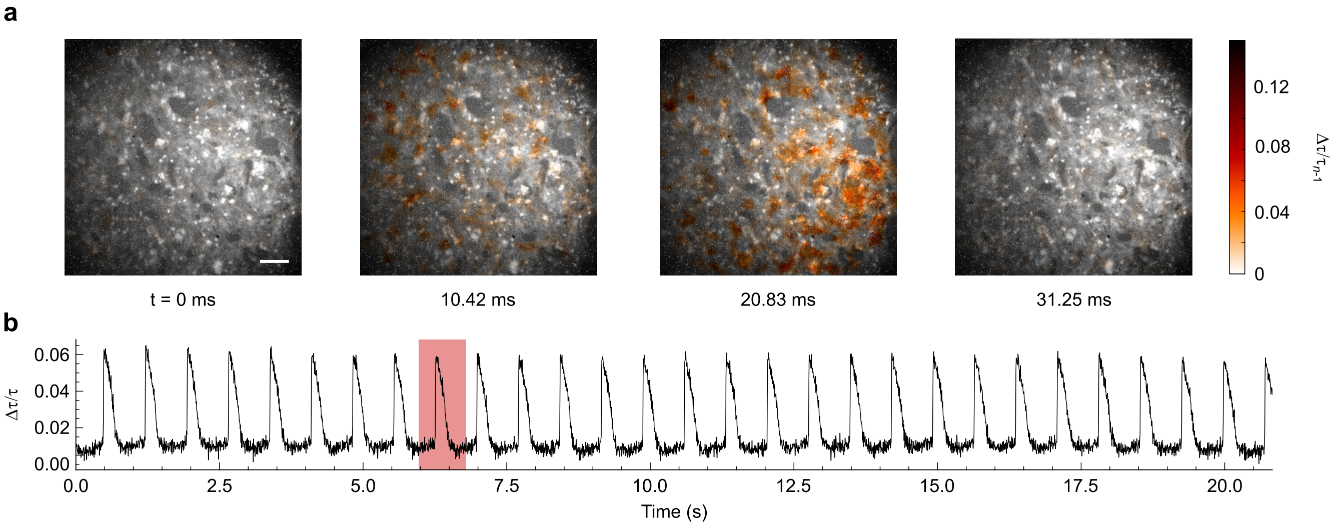


Supplementary Fig. 5. **Widefield fluorescence lifetime imaging of action potential propagation in a hiPSC-CM monolayer.** (**a**) $\Delta\tau/\tau_{n-1}$ time-lapse frames showing the propagation of the action potential wavefront in the hiPSC-CM monolayer. White scale bar, 100 µm. (**b**) Action potential time trace recorded over 20 s with the beat shown in (**a**) highlighted in red.

signal. Figure 4a shows the evolution of $\Delta\tau/\tau_{n-1}$ across the field of view during sequential beats. The activation maps derived from the fluorescence lifetime and from intensity (Fig. 4b) demonstrate that both modalities capture the same spatiotemporal propagation patterns of calcium activation throughout the tissue. As expected, the signal-to-noise ratio of the lifetime traces is lower than that of intensity, but the essential features of wave propagation remain accessible, confirming that the FLIM signal can be used for quantitative mapping of calcium activation dynamics.

#### WIDEFIELD IMAGING OF ACTION POTENTIAL PROPAGATION

In order to showcase our system's ability to spatially resolve action potential (AP) wave propagation in a hiPSC-CM monolayer stained with FluoVolt and imaged across an $816 \times 816$ µm FOV. Supplementary Fig. 5a illustrates the spatial mapping of AP propagation captured with our system with the corresponding AP time trace shown in Supplementary Fig. 5b. In this experiment, we applied our temporal cleaning process to enhance the clarity of the recorded signals. The refined recordings allow us to visualize the sequence of activation with high fidelity.

Supplementary Fig. 5a shows a series of time-lapse frames that map the progression of the action potential wavefront ($\Delta\tau/\tau_{n-1}$). Each frame has been processed to minimize temporal noise, as stated above. These detailed visualizations confirm the ability of our system to capture rapid electrophysiological events.

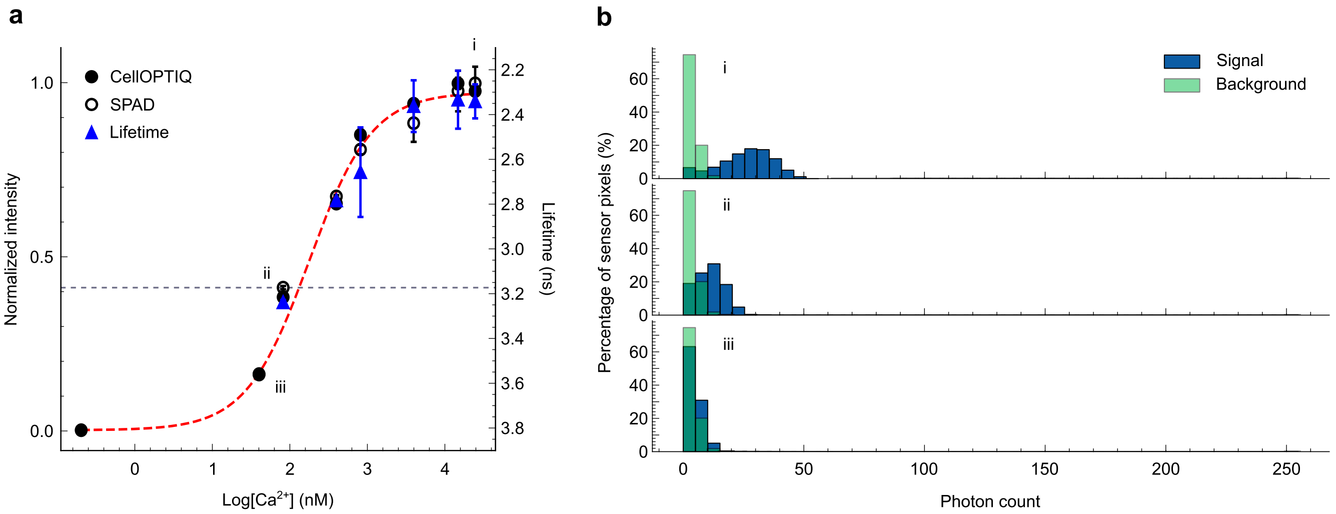


Supplementary Fig. 6. **Calibration curves relating fluorescence emission to calcium ion concentration and SPAD noise analysis.** (**a**) Normalized fluorescence intensity of Cal-520, AM measured with both CellOPTIQ system (solid circles) and the SPAD array (open circles) at varying free $Ca^{2+}$ concentrations. The sigmoidal curve is fitted to the CellOPTIQ data but shows good agreement with both imaging systems. Fluorescence lifetime values (blue triangles) are also plotted down to a threshold (gray dashed line) below which noise dominates and FLIM measurements are no longer reliable. Error bars represent the standard deviation. (**b**) Histograms of photon counts captured by the SPAD for the three points highlighted in (**a**). Below log[$Ca^{2+}$] values of 1.9 nM (iii) the distribution of the noise (green) becomes indistinguishable from that of the signal (blue). This overlap limits the ability to accurately extract calcium-dependent fluorescence lifetime changes at very low ion concentrations, imposing a lower threshold on the measurements in (**a**).

#### CALIBRATION OF FLUORESCENCE LIFETIME TO CALCIUM ION CONCENTRATION

Here we show the calibration curve used to convert fluorescence lifetime data into quantitative intracellular $Ca^{2+}$ concentrations. A series of calibration solutions were generated by mixing Ca-EGTA and EGTA buffers with 1 µM Cal-520, potassium salt, yielding free $Ca^{2+}$ levels ranging from 0.2 nM to 25 µM. A 200 µL aliquot of each solution was dispensed into a 96-well glass bottom plate and imaged using our FLIM system and a commercially available CellOPTIQ system (Clyde Biosciences, considered a gold standard for cardiac electrophysiology).

First, to ensure that the measurements from our system agree with standard calibration practices, we measured the fluorescence intensity of each well on a CellOPTIQ system to obtain the standard sigmoidal calibration curve reported for Cal-520, AM [3]. We then repeated this measurement on our system using the time-integrated intensity images from the SPAD as a comparison. These two data are shown in Supplementary Fig. 6a with the sigmoidal fit generated for the CellOPTIQ data. This fit gives a dissociation constant ($K_d$) of $186 \pm 3$ nM. The strong agreement between these data points confirms that the samples behaved similarly in both imaging systems for the full duration of the experiment. We then evaluated the average fluorescence lifetime of each well using the two gated images output by the SPAD, captured synchronously with the SPAD intensity data. In other words, the lifetime values blue triangles in Supplementary Fig. 6a) correspond directly to the well-documented intensity behavior of the dye (black circles).

Supplementary Fig. 6b shows histograms of the intensity data (photon counts) measured by the SPAD at three different free calcium concentrations – log[$Ca^{2+}$] = 4.4, 1.9 and 1.6 nM. At the lowest concentration (1.6 nM), the distribution of signal photon counts shows a complete overlap with the distribution of the background noise, making it impossible to distinguish noisy pixels from those containing a true signal. This sets a practical lower threshold for FLIM-based measurements, indicated in Supplementary Fig. 6a with a gray dashed line. Any estimate of fluorescence lifetimes below this threshold would be inaccurate and dominated by noise. Importantly, our system still provides accurate lifetime values in physiologically relevant calcium ranges.

To ensure consistency between FLIM and intensity-based calibrations, we also used the sigmoidal fit derived from the CellOPTIQ intensity data to estimate the fluorescence lifetime behavior across all calcium concentrations. Using the dissociation constant ($K_d = 186$ nM), the mean measured lifetime at maximal calcium ($\tau_{\text{max.}} = 2.41$ ns) and the interpolated lifetime at $K_d$ ($\tau = 3.10$ ns), we calculated the lifetime of the calcium-free form of the dye as $\tau_{\text{min.}} = 3.76$ ns. These values define a sigmoidal relationship between fluorescence lifetime and calcium concentration, modeled by the equation

$$[\text{Ca}^{2+}] = K_d \cdot \frac{\tau - \tau_{\text{min.}}}{\tau_{\text{max.}} - \tau}. \tag{3}$$

This overlapping calibration curve confirms the strong correspondence between intensity- and lifetime-based mea-

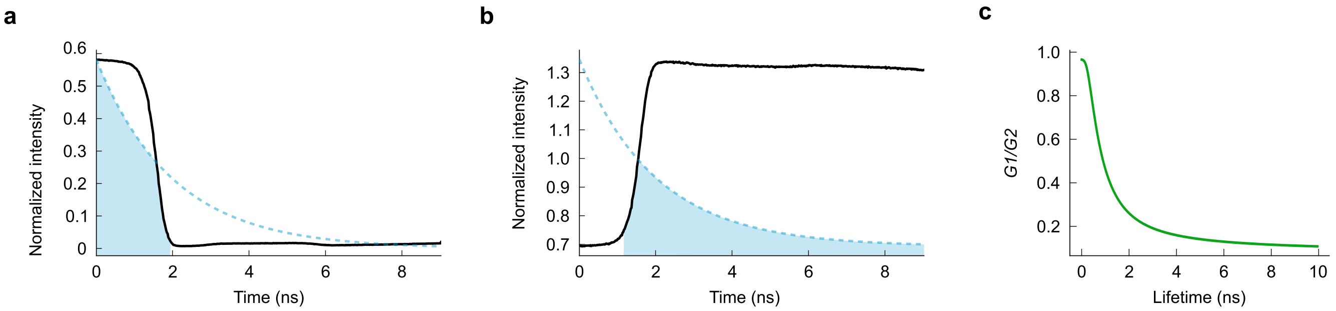


Supplementary Fig. 7. **Gate profiles and lookup table for rapid lifetime determination.** (**a**, **b**) Gate (**a**) and antigate (**b**) profiles from the SPAD overlaid with simulated fluorescence decay for visualization of data capture. The shaded regions represent the different phases of the fluorescence decay captured by each channel. (**c**) Example lookup table relating the ratio between gate and antigate channels with fluorescence lifetime.

surements, providing a robust method for converting FLIM data to intracellular calcium concentrations. These results validate the performance of our FLIM system for calcium quantification and confirm that Cal-520, AM exhibits consistent, well-characterized behavior across both imaging platforms when measured in solution. For all intracellular measurements presented in the main manuscript, we calibrated our time-course data using the established intracellular $K_d$ for Cal-520, AM (320 nM) [3], which accounts for the altered binding affinity of the dye in intracellular environments.

#### RAPID LIFETIME DETERMINATION GATING

In Supplementary Fig. 7 we show examples of the gate and antigate profiles of the SwissSPAD3 camera. Supplementary Fig. 7a, b show examples of the fixed gate positions used with simulated fluorescence decays to demonstrate the data capture paradigm. Each gate counts the number of photons arriving during the duration it remains open, illustrated by the highlighted regions under the blue decay curves.

The accuracy of the RLD approach depends on the precise timing of the gate relative to the fluorescence decay, with the highest accuracy coming from gating up to approximately the expected lifetime. To enable rapid lifetime estimation, a lookup table is used to map the measured gate-to-antigate ratio ($G1/G2$) directly to the fluorescence lifetime, as illustrated in Supplementary Fig. 7c. This calibration captures the expected ratio-lifetime relationship for the example gate configuration shown in Supplementary Fig. 7a, b. The calibration of the lookup table is most effective when the ratio-lifetime curve has a steep slope in the expected lifetime range, so that even small changes in the measured ratio correspond to clear changes in lifetime. This slope and its position are optimized by approximately matching the gate duration to the anticipated lifetime.

#### POISSONIAN NOISE STATISTICS IN INTENSITY AND LIFETIME MEASUREMENTS

To illustrate how Poissonian noise in photon counts propagates into the distribution of lifetimes estimated by the RLD method, we performed 1,000 Monte Carlo simulations of monoexponential decay curves at three nominal lifetimes ($\tau_{\mathrm{sim.}} = 2.5$, 5.0 and 7.5 ns). Photon arrival times were binned into two gates (experimentally measured for our SPAD and averaged across the sensor area) and sampled according to a Poisson process with a fixed noise level. Histograms of the summed counts across both gates (Supplementary Fig. 8a–c) demonstrate the intrinsic shot-noise–limited variance of the Poisson distribution under identical conditions.

When these noisy counts are converted to lifetime estimates using the dual-gated RLD method outlined above, the resulting lifetime distributions (Supplementary Fig. 8d–f) show the same Poissonian statistics. For $\tau = 2.5$ ns (d), the estimate distribution remains approximately Gaussian, while for $\tau = 5.0$ ns (e) the spread increases, and for $\tau = 7.5$ ns (f) the deviation of the lifetime from the fixed gate interval further amplifies variability (manifested as an asymmetric tail). This transfer of count variation into lifetime uncertainty explains the increased variation in $\tau$ observed in our calcium calibration measurements (Supplementary Fig. 6) under shot-noise limited conditions.

The lookup table is generated to be optimized for the experimental data with a gate width of $\sim 2$ ns, such that using it for longer lifetimes as seen here would be impractical. This is illustrated by the increasing standard deviation on $\tau_{\mathrm{est.}}$ shown in Supplementary Fig. 8d-f. If one wished to examine longer lifetime samples experimentally, a different gate width should be chosen and a new lookup table generated.

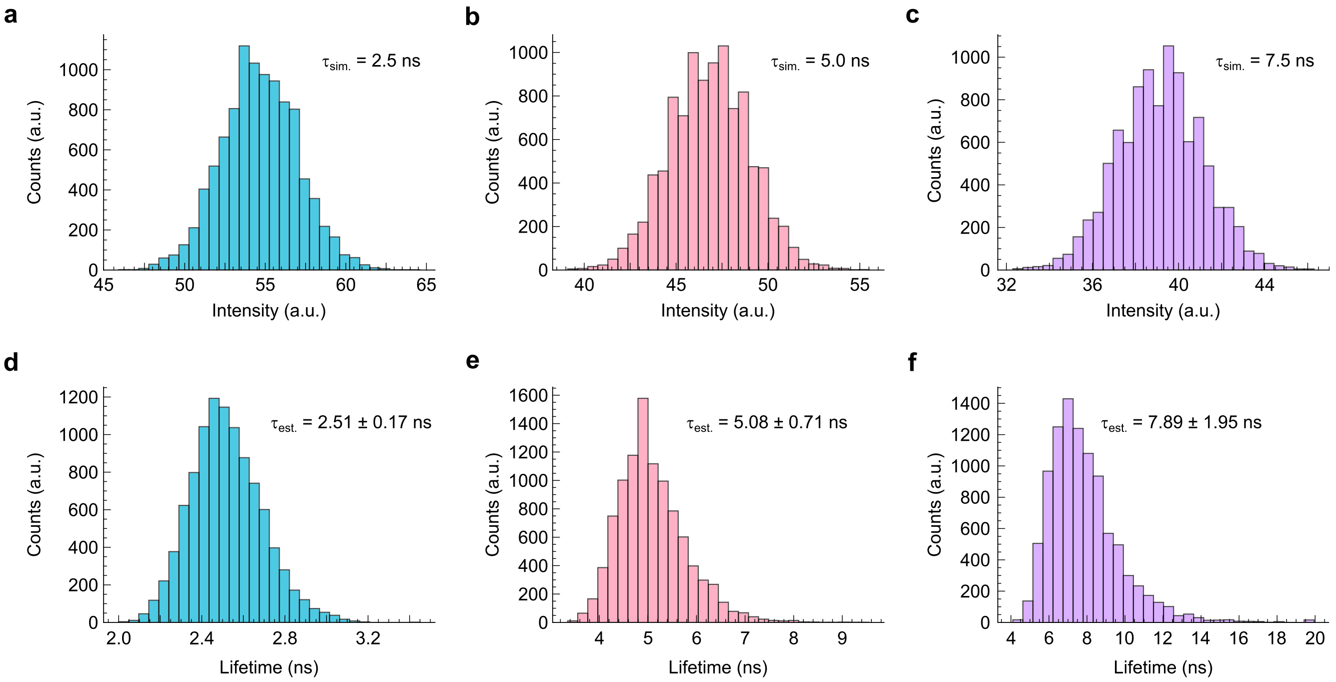


Supplementary Fig. 8. **Impact of Poissonian noise on dual-gated lifetime estimates.** Histogram of total photon counts (gate+antigate) over 1,000 simulations for (**a**) $\tau$ =2.5 ns, (**b**) $\tau$ = 5.0 ns and (**c**) $\tau$ =7.5 ns. (**d**–**f**) Corresponding histograms of lifetime estimates from the dual-gated RLD method: (**d**) $\tau$ = 2.5,ns, (**e**) $\tau$ = 5.0,ns, (**f**) $\tau$ = 7.5,ns. The increasing spread and asymmetric tail at longer $\tau$ reflect how this method begins to fail when the detected lifetime deviates greatly from the fixed gate width ($\sim$ 2 ns here).

#### TEMPORAL CLEANING

Here, we show pictorially the steps of the temporal cleaning procedure, used to suppress fluctuations in three-dimensional spatially resolved data cubes (originally reported in [4]). Due to frame-to-frame variations in the number of photons collected, the pixel-wise lifetime calculated will fluctuate from frame-to-frame over the course of a video. When plotting the instantaneous change in wavefront position ($\Delta\tau/\tau_{n-1}$) or traditional comparisons to a baseline frame ($\Delta\tau/\tau_0$), these fluctuations will present more obviously and obscure the desired signal with noise, as seen in Supplementary Fig. 9a. By isolating a single pixel (Supplementary Fig. 9b) and plotting its time-trace during physiological activity (Supplementary Fig. 9c) we see the effect of these rapid fluctuations. By fitting an analytical expression to this waveform, we can effectively remove the noise and completely suppress variations from pixels where electrical activity is not observed. To approximate both calcium and action potential waves, we fit a skew-normal distribution of the form

$$f(x) = 2\phi(x)\,\Phi(\alpha x)\,, \tag{4}$$

where

$$\phi(x) = \frac{1}{\sqrt{2\pi}} e^{-\frac{x^2}{2}}\,, \tag{5a}$$

$$\Phi(x) = \int_{-\infty}^{x} \phi(t)\,\mathrm{d}t = \frac{1}{2}\left[1 + \mathrm{erf}\left(\frac{x}{\sqrt{2}}\right)\right]. \tag{5b}$$

The shape of the distribution is controlled by $\alpha$, with $\phi(x)$ and $\Phi(x)$ representing the standard normal probability density and the cumulative distribution functions, respectively. This fitted data cube can then be visualized (Supplementary Fig. 9d) to show spatially resolved fluorescence lifetime changes devoid of noise fluctuations.

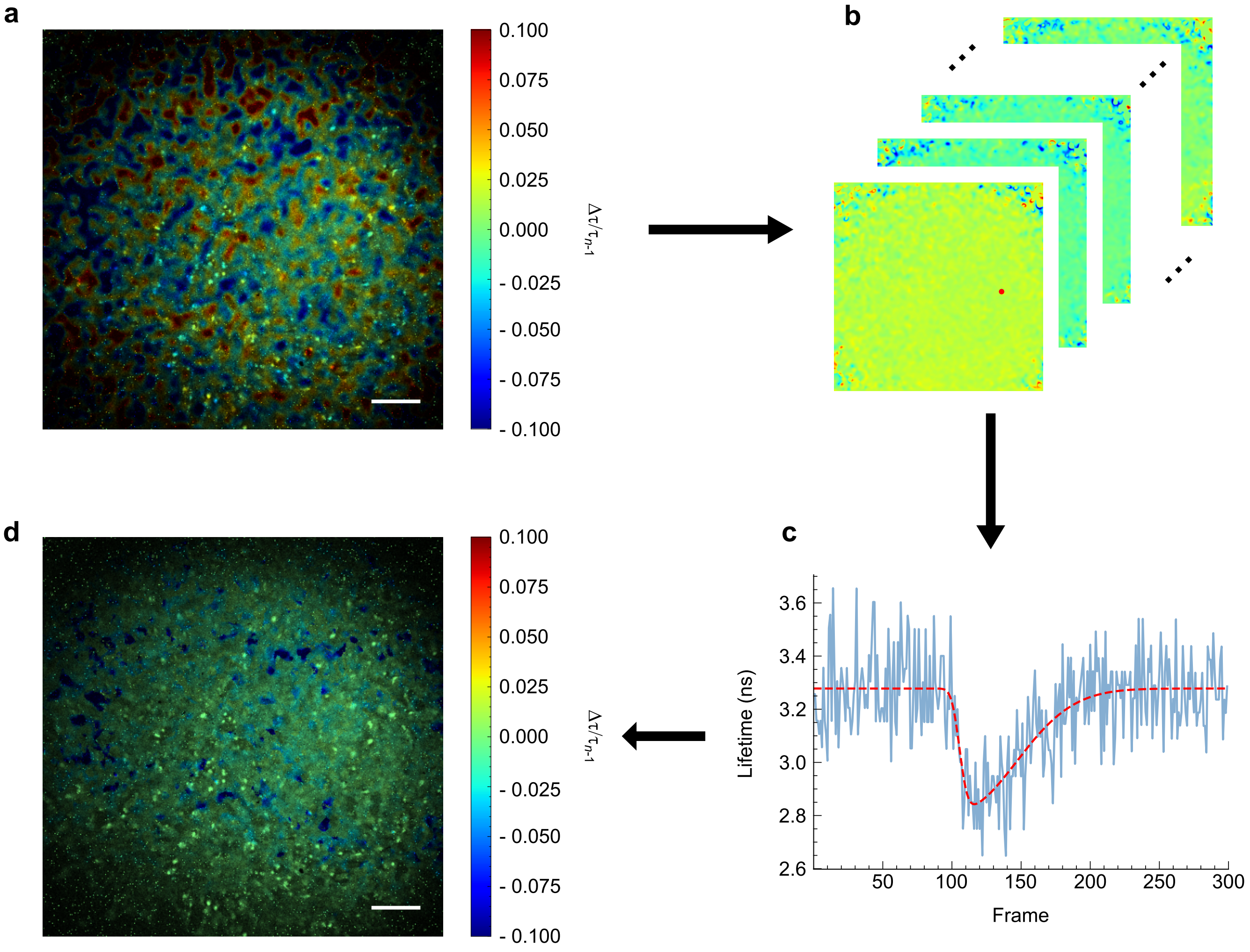


Supplementary Fig. 9. **Temporal cleaning of three-dimensional data cube.** (**a**) $\Delta\tau/\tau_{n-1}$ image from uncleaned FLIM data. (**b**) Raw FLIM frames. (**c**) Single pixel (red in (**b**)) showing calcium wave and fitted curve. (**d**) Spatially cleaned frame showing $\Delta\tau/\tau_{n-1}$ wavefront (blue) with no noise. White scale bar, 200 µm.


[1] L. Sala, B. J. van Meer, L. G. Tertoolen, J. Bakkers, M. Bellin, R. P. Davis, C. Denning, M. A. Dieben, T. Eschenhagen, E. Giacomelli, C. Grandela, A. Hansen, E. R. Holman, M. R. Jongbloed, S. M. Kamel, C. D. Koopman, Q. Lachaud, I. Mannhardt, M. P. Mol, D. Mosqueira, V. V. Orlova, R. Passier, M. C. Ribeiro, U. Saleem, G. L. Smith, F. L. Burton, and C. L. Mummery, MUSCLEMOTION, Circulation Research **122**, e5 (2018).

[2] F. Cerignoli, D. Charlot, R. Whittaker, R. Ingermanson, P. Gehalot, A. Savchenko, D. J. Gallacher, R. Towart, J. H. Price, P. M. McDonough, and M. Mercola, High throughput measurement of $Ca^{2+}$ dynamics for drug risk assessment in human stem cell-derived cardiomyocytes by kinetic image cytometry, Journal of Pharmacological and Toxicological Methods **66**, 246 (2012).

[3] AAT Bioquest, Inc., Cal-520®, AM Product Datasheet, `https://www.aatbio.com/products/cal-520-am` (2025), accessed: 2025-04-24.

[4] G. Gariepy, N. Krstajic, R. Henderson, C. Li, R. R. Thomson, G. S. Buller, B. Heshmat, R. Raskar, J. Leach, and D. Faccio, Single-photon sensitive light-in-fight imaging, Nature Communications 2015 6:1 **6**, 1 (2015).
